# CD3e-immunotoxin spares CD62L^lo^ Tregs and reshapes organ-specific T-cell composition by preferentially depleting CD3e^hi^ T cells

**DOI:** 10.1101/2022.07.25.501205

**Authors:** Shihyoung Kim, Rajni Kant Shukla, Hannah Yu, Alice Baek, Sophie G Cressman, Sarah Golconda, Ga-Eun Lee, Hyewon Choi, John C Reneau, Zhirui Wang, Christene A. Huang, Namal P. M. Liyanage, Sanggu Kim

**Affiliations:** Department of Veterinary Biosciences, The Ohio State University, Columbus, OH 43210, USA; Department of Microbial Immunity and Infection, The Ohio State University, Columbus, OH 43210, USA; The Ohio State University, Division of Hematology, Columbus, OH 43210, USA; Department of Surgery, University of Colorado Denver Anschutz Medical Campus, 12700 East 19^th^ Ave, Aurora, CO, 80045, USA; Infectious Disease Institute, The Ohio State University, Columbus, OH 43210, USA

**Keywords:** CD3e immunotoxin, tissue-resident T cells, regulatory T cells, tolerance induction, T-cell lymphoma therapy, immunotherapy, CD3e expression, organ-to-organ variation

## Abstract

CD3-epsilon(CD3e) immunotoxins (IT), a promising precision reagent for various clinical conditions requiring effective depletion of T cells, often shows limited treatment efficacy for largely unknown reasons. Tissue-resident T cells that persist in peripheral tissues have been shown to play pivotal roles in local and systemic immunity, as well as transplant rejection, autoimmunity and cancers. The impact of CD3e-IT treatment on these local cells, however, remains poorly understood. Here, using a new murine testing model, we demonstrate a substantial enrichment of tissue-resident Foxp3+ Tregs following CD3e-IT treatment. Differential surface expression of CD3e among T-cell subsets appears to be a main driver of Treg enrichment in CD3e-IT treatment. The surviving Tregs in CD3e-IT-treated mice were mostly the CD3e^dim^CD62L^lo^ effector phenotype, but the levels of this phenotype markedly varied among different lymphoid and nonlymphoid organs. We also found notable variations in surface CD3e levels among tissue-resident T cells of different organs, and these variations drive CD3e-IT to uniquely reshape T-cell compositions in local organs. The functions of organs and anatomic locations (lymph nodes) also affected the efficacy of CD3e-IT. The multi-organ pharmacodynamics of CD3e-IT and potential treatment resistance mechanisms identified in this study may generate new opportunities to further improve this promising treatment.

## Introduction

CD3e-IT, a fusion of the catalytic domain of diphtheria toxins with the CD3e-binding portion of a CD3e antibody is a promising precision medicine for an effective T-cell depletion. CD3e-IT has shown preclinical and clinical efficacy for the treatment of T-cell lymphomas,(1) autoimmune diseases(2) and transplant rejection,(3– 7) and is currently under evaluation as a preconditioning regimen for cell therapy. A recent *Resimmune*® [A-dmDT390-bisFv(UCHT1)](8) clinical trial showed notable response rates of 47–74% in patients with intermediate-stage (stage-IB/IIB or mSWAT score < 50) cutaneous T-cell lymphomas (CTCL).(9) Its effects on stage III/IV CTCL and peripheral blood T-cell lymphomas, however, seemed limited for reasons that remain unclear. The limited efficacy of immunotoxins has been attributed to several potential factors, including pre-existing antibodies or immunogenicity against CD3e-ITs, the lower levels of CD3e or insensitivity of lymphoma cells in advanced CTCL, the lack of penetration of the immunotoxins into target cell sites, and protective tumor microenvironment.(9–13) The impact of pre-existing antibodies or immunogenicity against the “second generation” recombinant CD3e-IT seemed to be mild and insignificant.(14–16) Other factors that may influence CD3e-IT efficacy include the differential expression of the TCR/CD3 antigen receptor complexes,(17, 18) the CD3e isoforms in the TCR/CD3 complexes(19, 20), the polymorphism of CD3e(21), and other cell-intrinsic factors of basically every step of toxin-mediated cell-killing processes(22). The impact of these cell-intrinsic and local factors on *in vivo* CD3e-IT efficacy remains unclear.

CD3e-IT has also proven effective in inducing long-term allograft acceptance in nonhuman primates and swine models(3–7, 23–29). Mounting evidence points to the critical roles of regulatory T cells (Tregs) in inducing tolerance in various treatment settings,(30–33) but their roles in CD3e-IT-mediated tolerance remain unclear. CD3e-IT has shown to be more effective in depleting T cells in the lymph nodes (LN) and skin than antibodies(34, 35) and reliably prolonged allograft survival.(36, 37) CD3e-IT alone or in combination with a short-term immunosuppressive chemotherapy, however, often showed limitations in sustaining graft function or survival.(36, 38–40) Recent CD3e-IT-mediated donor hematopoietic stem cell transplant studies have demonstrated promising results where long-term acceptance of allografts seemed to be finally achievable through mixed chimerism in monkey and swine models.(4, 5, 29) Unlike the similar studies utilizing anti-T-cell antibodies,(41–43) there was however no clear evidence that T-cell anergy or regulation by Tregs serve as the dominant mechanism of allograft tolerance. The CD3e-IT-mediated transient chimerism, instead, facilitated the robust and stable humoral immune modulation of donor-specific antibody responses.(44) Treg infiltration into an allograft has been reported in a miniature swine model, suggesting a potential local regulatory component.(4) Nevertheless, the survival and roles of Tregs in CD3e-IT-mediated tolerance induction remain unclear.

This study presents a detailed pharmacodynamic evaluation of CD3e-IT treatment with a particular focus on the survival of Tregs in different organs. Tissue-resident T cells rarely circulate and instead remain stably parked in the tissue parenchyma of local organs.(45–48) These local T cells collectively account for the largest T-cell subset in the body. They have been shown to participate in protection from infection and cancer and are associated with allergy, autoimmunity, inflammatory diseases and transplant rejection.(45) Our understanding of these local cells in CD3e-IT treatment settings remains surprisingly poor. Tregs must home to LNs to establish antigen-specific T-cell tolerance.(49–51) The activated effector Tregs in LNs may then exert their suppressive functions both locally and systemically. Local Tregs showed a marked skewing of T-cell receptor (TCR) usage by anatomical location which contribute to shaping the unique peripheral Treg population in different secondary lymphoid organs.(52) Follicular T regulatory cells (Tfr) that primarily reside in B-cell follicles are critical in shaping humoral immune responses by controlling follicular T helper cells (Tfh) and B cells in the germinal center.(53, 54) Like Tfr, other tissue-resident Tregs in nonlymphoid organs or in allografts also play pivotal roles in organ-specific functions and homeostasis.(55) Treg infiltration into the graft also coincided with the metastable tolerance achieved in animals and humans.(38, 56, 57) Nevertheless, the impact of CD3e-IT on local tissue-resident Tregs or T cells in general remain largely uncharacterized. Only a few recent studies, including our own, have shown differential depletion efficiencies for tissue-resident T cells by anti-T-cell mAb and CD3e-IT treatments in a few selected organs.(35, 46) Most past studies have focused almost exclusively on total peripheral blood, lymph node and bone marrow cells in animal models(34, 37, 58–60) and total peripheral blood in humans,(1) without drawing any clear distinction between circulating and tissue-resident populations.

Despite the numerous advantages over other animal models, mouse models have been used only rarely for CD3e-IT studies, primarily due to the limited efficacy and toxicity of first-generation murine CD3e-ITs.(61) We have recently developed a new murine-version CD3e-IT (S-CD3e-IT) whose safety and treatment efficacy profile is comparable to those of the “second-generation” recombinant CD3e-ITs.(46) Here, we demonstrate the significant enrichment of Forkhead box transcription factor (Foxp3) Tregs in the murine model following CD3e-IT. We used the new murine testing model to systemically characterize Foxp3+ Treg enrichment in multiple organs – including the peripheral blood, spleen, lung, Peyer’s Patches, 5 types of lymph nodes (mesenteric, inguinal, mandibular, mediastinal, and lumbar), thymus, and bone marrow – using intravascular staining techniques to analyze circulating and tissue-resident cells separately.(46–48) We found S-CD3e-IT effectively deplete both circulating and tissue-resident T cells, but organ-to-organ variations were evident. S-CD3e-IT preferentially depleted CD4+ T-cell subsets that express high levels of CD3e (CD3e^hi^), sparing T-cells with low levels of CD3e (CD3e^dim^). Differential surface expression of CD3e molecules among T-cell subsets, as well as organ-to-organ variations in CD3e expression, drive Treg enrichment and reshape organ-specific T-cell composition. As a result, effector Tregs (CD62L^lo^) with the CD3e^dim^ phenotype were substantially enriched in the tissue-resident pools of secondary and nonlymphoid organs.

## Results

### New murine CD3e-IT effectively depletes tissue T cells with notable organ-to-organ variations

For a detailed pharmacodynamics analysis of CD3e-IT, we employed a new murine testing system that we developed recently(Figure 1A).(46) Anti-murine-CD3e-ITs (S-CD3e-ITs) were generated by conjugating streptavidin-saporin (Advanced Technology System) with non-mitogenic CD3e-mAb (145-2C11 with Fc-silent™ murine IgG1, Absolute Antibody) as described previously.(46) S-CD3e-IT was highly precise and efficacious in depleting both circulating and tissue-resident T cells in mice (Figure 1B). We tested S-CD3e-IT in both nonimmunized and OVA-immunized C57BL/6J mice. A systemic immunization induces T-cell activation and infiltration into uninflamed tissues, including both secondary and nonlymphoid organs.(45, 62– 65) Akin to previous reports,(66) the LN cells of OVA-immunized mice showed a notable increase in surface expression of CD69 (a marker of tissue residency) on CD4+ T cells compared to those in nonimmunized mice (Supplemental Figure 1A).

**Figure 1.**
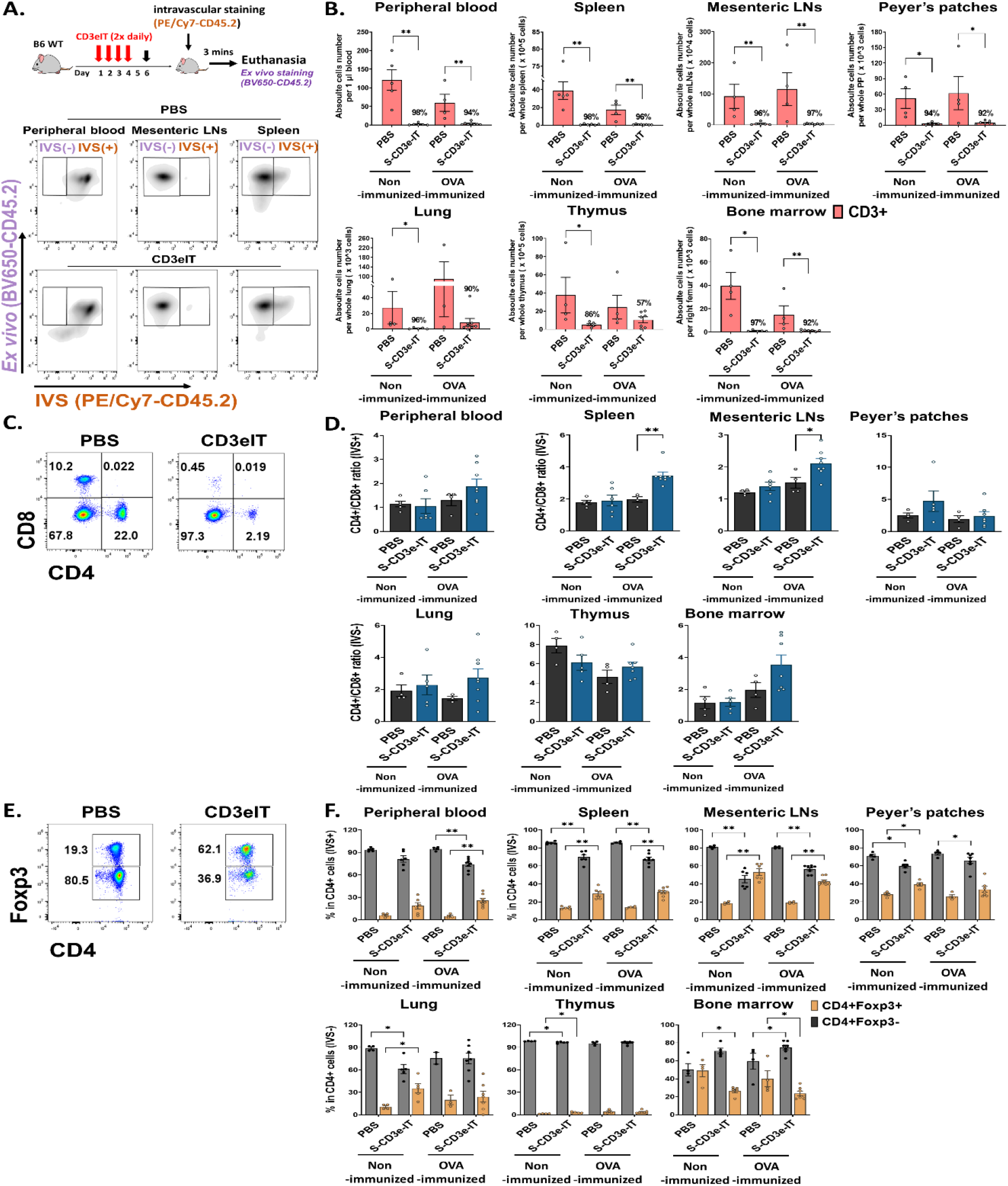
CD4+Foxp3+ Tregs cells enrichment following S-CD3e-IT treatment. (A) The diagram shows an S-CD3e-IT treatment and intravascular staining strategy to analyze the effect of S-CD3e-IT efficacy on different Tissue-resident T cells The flow cytometry gating strategy shows circulating (IVS+) and tissue-resident cells (IVS-) separation in peripheral blood, mesenteric LNs, and spleen. (B) Absolute number of CD3+ cells (y-axis) is shown for the peripheral blood (IVS+) and tissue-resident (IVS-) portions of spleen, mesenteric LNs, Peyer’s patches, lung, thymus, and bone marrow. Absolute cell numbers were calculated based on CountBright Absolute counting beads and total cell counts per organ. (C-D) Flow cytometry for CD4+ and CD8+ T cells (C) and the ratios of CD4+ to CD8+ for all tested organs (D) are shown. (E-F) Flow cytometry for CD4+Foxp3- and CD4+Foxp3+ T cells (E) and % of these cells in CD4+ cells (F) are shown for all tested organs. Nonimmunized mice treated with PBS (*n* = 4∼5, depending on organs), or S-CD3e-IT (*n* = 4∼6), OVA-immunized mice treated with PBS (*n* = 3∼4), or S-CD3e-IT (*n* = 7∼8), were compared. (* *p* < 0.05 and ** *p* < 0.01).

Both nonimmunized and OVA-immunized C57BL/6J mice were then treated with S-CD3e-IT (15μg twice daily by retro-orbital injection) for 4 consecutive days, as described previously.(46) On Day 6, these mice were subjected to intravascular staining [IVS; by retro-orbital injection of anti-CD45.2 mAb (PE/Cy7-CD45.2)] followed by euthanasia 3 minutes later. Total organ cells were then *ex vivo* stained with an antibody pool containing another anti-CD45.2 mAb (BV650-CD45.2) (Figure 1A). The circulating cells in the vasculature were thus identified as IVS+ and the tissue-resident cells in the tissue parenchyma at the time of injection as IVS−. As expected, in all PBS-treated groups, >99% of peripheral blood CD45+ cells were IVS+, whereas about 98% of LN- and thymus-isolated cells were tissue-resident (IVS−;Supplemental Figure 2).(47) Consistent with previous reports,(46, 47) >20% of lung-isolated cells and 70-80% of spleen- and BM-isolated cells were IVS− (Supplemental Figure 2).

T-cell depletion rates were >90% for both IVS+ and IVS-pools of all tested organs (except for the thymus and a few local LNs), with notable organ-to-organ variations (Figure 1B and Supplemental Figure 3A). The low depletion rates in the thymus may reflect the functional properties of the thymus which was continuing to produce new CD3+ T cells in these mice (Supplemental Figure 4). Different LN types showed highly variable depletion rates depending on the anatomic sites of LNs (discussed below in more detail; see Fig.5). Both CD4+ and CD8+ cells were effectively depleted. Unlike the reduced CD4/CD8 ratios in CD3-mAb treatment,(18) however, the CD4/CD8 ratios in CD3e-IT-treated mice were increased in most organs (Figure 1D and Supplemental Figure 3B), indicating preferential depletion of CD8+ T cells over CD4+ T cells by CD3e-IT.(40, 46) Although T-cell depletion rates were slightly lower in OVA-immunized mice, the organ-to-organ variations were generally consistent between the nonimmunized and OVA-immunized groups (Figure 1B).

### Tissue-resident CD4+ FoxP3+ Tregs were enriched in both secondary lymphoid and non-lymphoid organs

The fraction of Foxp3+ Tregs in PBS control mice moderately varied among secondary lymphoid and non-lymphoid organs, appearing in average 5.2% to 28.3% of CD4+ T cells (Figure 1F), consistent with previous reports in humans and mice.(67–71) S-CD3e-IT treatment significantly enriched Foxp3+ Tregs in the tissue-resident pool (IVS−) of most organs, but not in the bone marrow (Figure 1F and Supplemental Figure 3C). Bone marrow is a known reservoir for Tregs, maintaining strikingly high levels of local Tregs under homeostatic conditions by actively, rather than passively, retaining these cells via CXCL12/CXCR4 signals(69). The reduced bone marrow Treg fraction – from 40.1-49.0% (in PBS mice) to 23.8-26.7% (in S-CD3e-IT-treated mice) – may indicate some perturbation of the bone marrow’s ability to actively retain Tregs in its parenchyma (Figure 1F). All other organs, however, showed varying levels of the enrichment of CD4+ FoxP3+ Tregs in local tissue sites after S-CD3e-IT treatment. Notably, the Foxp3+ Tregs accounted for nearly half of the total CD4+ T cells survived in LNs (Supplemental Figure 3C), an anatomic site orchestrating Treg activities in local organs(52, 72).

### Preferential depletion of CD3^hi^ T cells drives enrichment of Tregs with the CD3e^dim^ phenotype

In In CD3e-mAb treatment settings, Tregs have been shown to have a survival advantage over other T-cell subsets, likely due to the relatively low surface expression of CD3e molecules on CD4+ FoxP3+ Tregs compared to those of other T-cell subsets.(17, 18) The TCR/CD3 complexes on the surface of Tregs are also shown to be enriched in CD3e isoforms with an undegraded N terminal, which probably further contributes to the resistance of these cells to CD3 antibody-mediated cell death.(19, 20) Consistent with these reports, CD4+Foxp3-T cells in PBS control mice showed the highest levels of CD3 expression on the cell surface, whereas CD4+Foxp3+ cells showed the lowest levels among tested T-cell subsets (Figure 2, A and B). Unlike CD3e-mAbs that work through TCR/CD3 modulation,(46, 73) however, the binding and internalization of CD3e-IT with CD3/TCR would likely kill the host cells instead of activating TCR/CD3 signals(46). Even a single molecule of ribosomal inactivating toxins, once internalized into a cell, can kill the cell by catalytically inhibiting protein synthesis.(74) We found that S-CD3e-IT treatment preferentially depleted T cells that display high levels of CD3e molecules (CD3e^hi^) on the cell surface – reflecting the controlled CD3e-IT dosage in our experiments – and spared T cells with a low-CD3e-expression phenotype (denoted as CD3^dim^) (Figure 2, A and B). The CD3^dim^ phenotype was clearly distinct from the CD3e modulation phenotype observed in the CD3e-mAb treatment; CD3e-mAbs nearly completely remove or internalize CD3e molecules on the surface of T cells without killing the cells (Figure 2A).(46, 73) CD8+ T cells display relatively lower CD3e molecules on the surface compared to CD4+ T cells; nevertheless, unlike CD3e-mAb treatment,(18) CD3e-IT preperentially depleted CD8+ T cells over CD4+ T cells.(40, 46)

**Figure 2.**
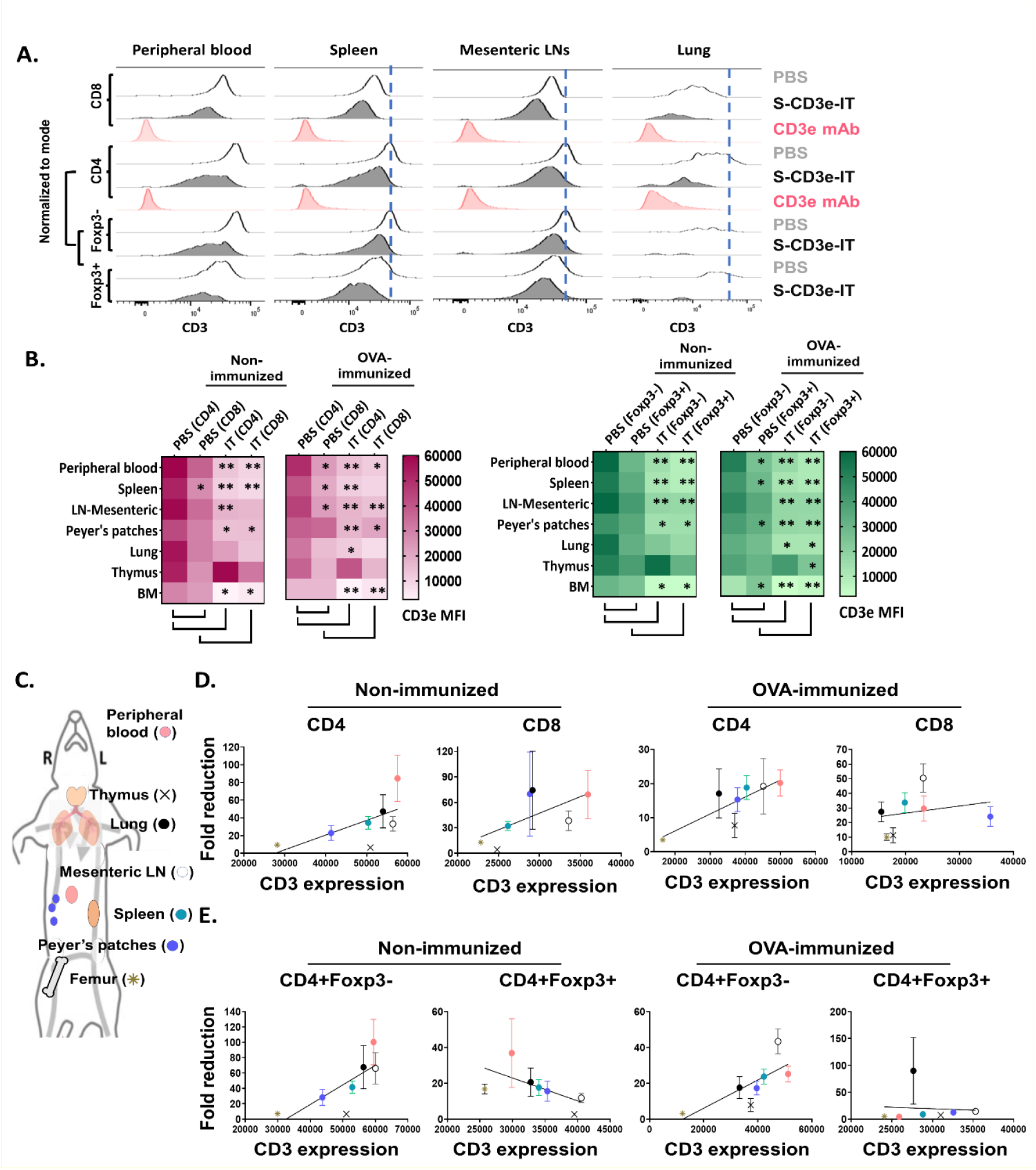
Differential CD3e on T cells drive an enrichment of CD3^dim^ Tregs with organ-to-organ variations. (A) CD3e expression (x-axis) on the CD8+, CD4+, CD4+Foxp3-, and CD4+Foxp3+ cells (y-axis, normalized to mode) are shown for the peripheral blood and tissue-resident portions (IVS-) of spleen, mesenteric LNs, and lungs. (B) Heatmaps show CD3e mean fluorescent intensities (MFI) for CD4+, CD8+, CD4+Foxp3- and CD4+Foxp3+ cells cells of all tested organs. Statistical comparison of CD3e MFI on two groups, CD4 (PBS) vs CD8 (PBS) and PBS-treated vs S-CD3e-IT-treated mice, is denoted with an asterisk (*) on the heat map. (* *p* < 0.05 and ** *p* < 0.01). (C) The diagram shows the symbol of all tested organs in (D) and (E). (D) A positive correlation between the fold reductions of CD4+ or CD8+ cell numbers in different organs after CD3e-IT treatment (y-axis) and CD3e MFI of these cells in PBS-treated animals (x-axis). CD4+ and CD8+ lymphocytes for the peripheral blood (red), and tissue-resident (IVS-) portions of spleen (green), mesenteric LNs (white), Peyer’s patches (purple), lung (black), thymus (cross), and bone marrow (asterisk) are compared. (E) A positive correlation between the fold reductions of CD4+Foxp3-cell numbers in different organs after CD3e-IT treatment (y-axis) and CD3e MFI of these cells in PBS-treated animals (x-axis). Note that this correlation was not evident for CD3+Foxp3+ T regs. Nonimmunized mice (PBS *n* = 4∼5, and S-CD3e-IT *n* = 4∼6, depending on organs); OVA-immunized mice (PBS *n* = 3∼4, and S-CD3e-IT *n* = 7∼8, depending on organs)

### Organ-to-organ variations in CD3e expression correlates to differential T-cell depletion rates indifferent organs

In PBS control mice, we also found notable organ-to-organ variations in the CD3e expression levels for each T-cell subset. Peripheral blood T cells and mesenteric LN-resident (IVS-) T cells consistently showed highest levels of CD3e mean fluorescent intensity (MFI) for nearly all T-cell subsets (except CD8+ T cells and Foxp3+ Tregs), whereas bone marrow-resident T cells consistently showed lowest (Figure 2, D and E). Notably, these differential CD3e MFI levels among tested organs were positively correlated with the depletion rates for CD4+ cells, CD8+ cells, and CD4+FoxP3-cells in these organs in S-CD3e-IT treated mice (Figure 2, D and E). However, such a pattern – reflecting the preferential depletion of T cells that display higher CD3e – was not evident with CD4+FoxP3+ Tregs (Figure 2E). This discrepancy suggests that – while CD3e-IT still preferentially depletes CD3e^hi^ Tregs over CD3e^lo^ Tregs – the organ-to-organ variations in the depletion of Tregs may be affected by other factors besides CD3e expression levels, including the intrinsic differences in the content of TCR/CD3 complexes (e.g. CD3e isoforms) and the relative insensitivity to external stimuli among Tregs in different organs.(17–20) Both the quantitative and qualitative variations in surface CD3e on T cell subsets, therefore, at least partially drive the observed variations in T-cell depletion and Treg enrichment in different organs after S-CD3e-IT treatment.

### CD62L^lo^ Tregs are enriched in secondary lymphoid and non-lymphoid organs

The L-selectin (CD62L) is a crucial lymphoid homing molecule. Treg populations can be largely classified as CD62L^hi^ and CD62L^lo^ Tregs. CD62L^hi^ Tregs are mostly naive and quiescent, whereas CD62L^lo^ Tregs are comprised of recently activated and short-lived cells.(75, 76) Both CD62L+ and CD62L-Tregs have shown a similar suppressive capacity for T-cell activation in vitro,(77, 78) but the CD62L+ Tregs – not the CD62L-counterparts – are likely the major population that provides long-lasting immune tolerance, protecting against lethal acute graft-versus-host disease(79, 80) and delaying diabetes in prediabetic nonobese diabetic mice.(78) Studies showed marked increase in CD62L+ Tregs in CD3e-mAb-treated mice.(81, 82) By contrast, in S-CD3e-IT-treated mice, we found that CD62L^hi^ Tregs were preferentially depleted by S-CD3e-IT in all test organs, except peripheral blood and bone marrow (Figure 3, A-C). As expected, the surviving Tregs were mostly the CD3^dim^ phenotype, except those in the thymus (Figure 3C). CD62L^hi^ Tregs (mostly CD44^lo^) also showed notably higher CD3e expression on the cell surface than CD62L^lo^ Tregs (mostly CD44^hi^) in PBS control mice (Figure 3C), consistent with a previous report that demonstrated a relatively higher CD3e expression on CD44-T cells than those of their CD44+ counterparts.(18)

**Figure 3.**
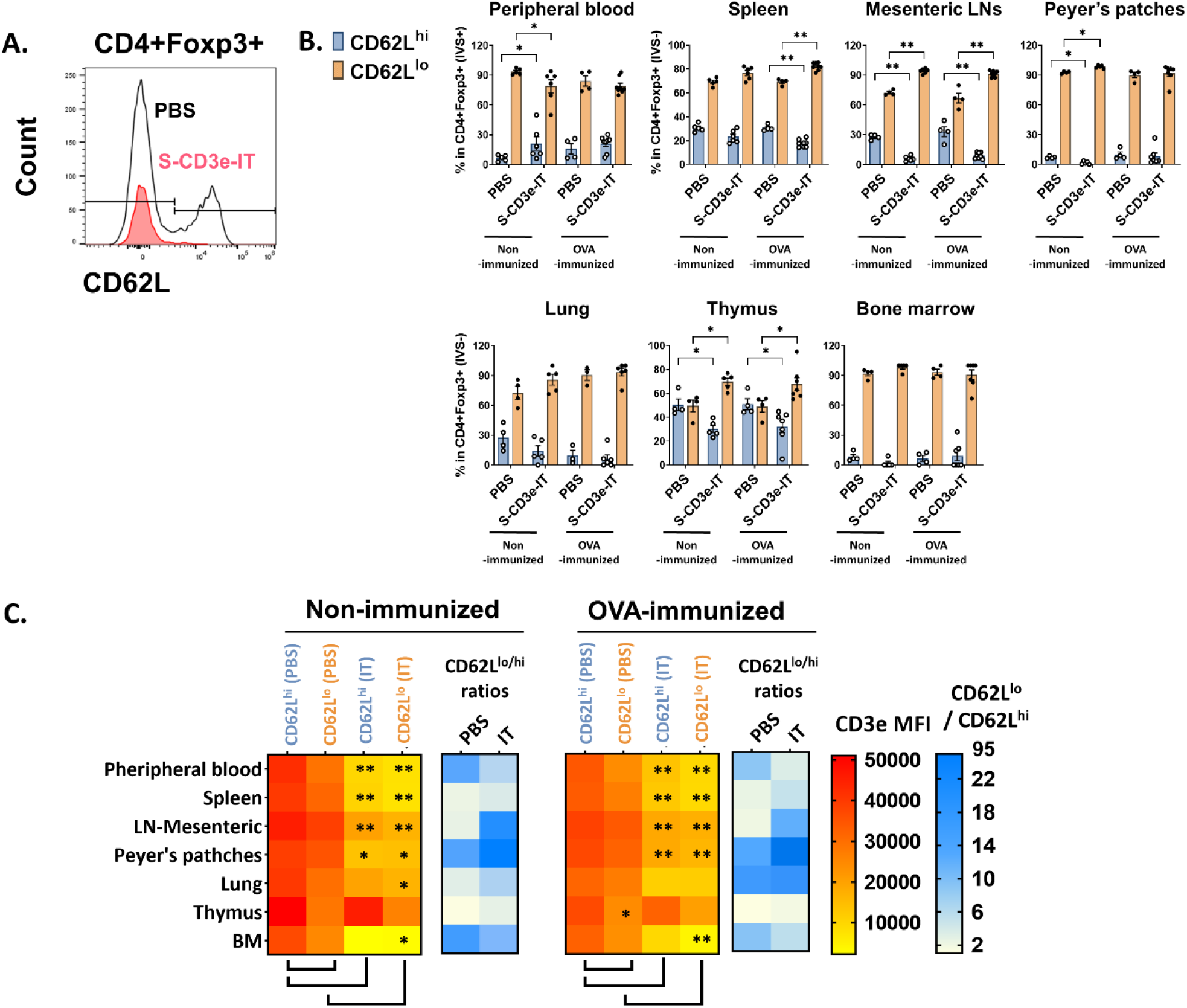
CD3^dim^CD62L^lo^ Treg enrichment correlates with CD3e expression. (A) The flow cytometry gating strategy for CD4+Foxp3+CD62L^lo^ and CD4+Foxp3+CD62L^hi^. (B) CD62L^lo^ and CD62L^hi^ (% in CD4+Foxp3+ cells) are shown for the peripheral blood (IVS+) and tissue-resident (IVS-) portions of spleen, mesenteric LNs, peyer’s patches, lungs, thymus, and bone marrow. (C) The heatmap of CD3e MFI (yellow to red) on CD4+Foxp3+CD62L^lo^ and CD4+Foxp3+CD62L^hi^ cells in tested organs. CD3e MFI on CD62L^lo^ was higher than CD62L^hi^ in all tested organs. A statistical comparison of CD3e MFI of two groups, CD62L^lo^ (PBS) vs CD62L^hi^ (PBS) and PBS-treated vs S-CD3e-IT-treated mice, is denoted with an asterisk (*) on the heat map. The heatmap of the CD62L^lo^-to-CD62L^hi^ ratios (white to blue). The ratio increased after S-CD3e-IT treatment except in peripheral blood and bone marrow. Nonimmunized mice were treated with PBS (*n* = 4∼5, depending on organs) or S-CD3e-IT (*n* = 4∼6). OVA-immunized mice were treated with PBS (*n* = 3∼4) or S-CD3e-IT (*n* = 7∼8). (* *p* < 0.05 and ** *p* < 0.01)

### Proliferation potentials of CD3e^dim^ CD4+ Foxp3+ T cells vs. their Foxp3-counterparts

CD3e-mAb treatment has yielded a transient increase in the relative fraction of Tregs – though not necessarily an increase in Tregs’ cell count – during the early stage of tolerance induction.(18, 73, 82–89) In CD3e-IT treated mice, the fraction of CD25+ Tregs were also significantly enriched within the tissue-resident (IVS−) pools of the secondary lymphoid and non-lymphoid organs (Figure 4A). Expression of CD25 (an α chain of the high-affinity IL-2 receptor) indicates potential sensitivity to IL-2 and lymphopenia-mediated induction, the primary modulators for the development and maintenance of Tregs.(90–92) Given that Tregs, particularly CD62L^lo^ effector Tregs, require TCR/CD3 activation to proliferate and maintain their functional properties.(93– 95) The activation and proliferation status of the surviving Tregs – mostly CD3e^dim^ CD62L^lo^ phenotype – were assessed by Ki67, CD44, CXCR5, and PD-1 expression. Our S-CD3e-IT treated mice demonstrated a marginal increase in Ki67 expression (a cell proliferation marker) for CD4+Foxp3+ Tregs in most organs (Figure 4B and Supplemental Figure 5). CD44, CXCR5, and PD-1 expression also showed only marginal increase in CD3e-IT-treated mice. By contrast, the CD4+Foxp3-counterparts showed a significant increase in CD44+ phenotypes and a 2-to 6-fold increase in Ki67 expression in peripheral blood and secondary lymphoid organs (Figure 4B and Supplemental Figure 5). Peripheral blood CD4+ Foxp3-T cells in OVA-immunized mice showed a >20-fold increase in PD-1 expression, a marker of exhausted T cells (Figure 4B and Supplemental Figure 4). Extensive homeostatic expansion of memory T cells has been a common event following CD3e-IT treatment(96).

**Figure 4.**
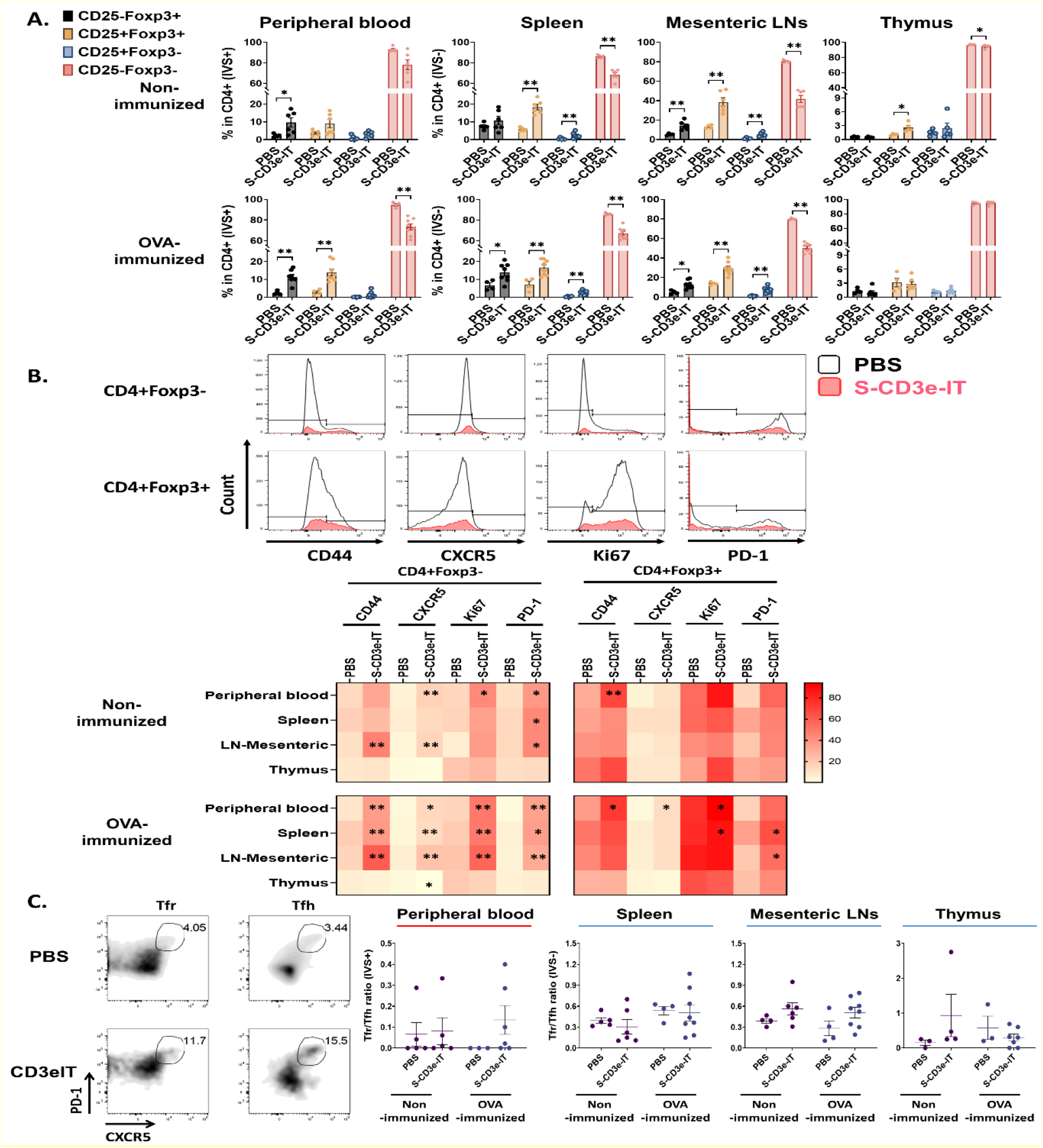
Changes in the functional phenotype of T cells after CD3e-IT treatment. (A) S-CD3e-IT treatment enriched CD25+ Tregs cells. CD25+ population (% in CD4+ cells) is shown for the peripheral blood and tissue-resident (IVS-) portions of spleen, mesenteric LNs, and thymus in PBS and S-CD3e-IT treated mice. (B) The flow gating strategy is shown to define CD44+, CXCR5+, Ki67+, and PD-1+ cells in CD4+Foxp3- and CD4+Foxp3+ cells. The heat map shows four functional phenotypic changes (light red to dark red) of CD4+Foxp3- and CD4+Foxp3+ T cells following S-CD3e-IT treatment. Statistical comparison between PBS-treated vs S-CD3e-IT-treated mice is denoted with an asterisk (*) on the Heat map. (C) The flow cytometry gating strategy is shown to define CXCR5+PD-1+ Tfr or Tfh. The CXCR5+PD-1+ Tfr-to-CXCR5+PD-1+ Tfh ratios are shown forperipheral blood and tissue-resident (IVS-) portions of the spleen, mesenteric LN, and thymus. Nonimmunized mice were treated with PBS (*n* = 4∼5, depending on organs) or S-CD3e-IT (*n* = 4∼6). OVA-immunized mice were treated with PBS (*n* = 3∼4) or S-CD3e-IT (*n* = 7∼8). (* *p* < 0.05 and ** *p* < 0.01).

### Notable increase in Tfr-to-Tfh ratios in mesenteric lymph nodes

We examined the Tfr-to-Tfh ratios in circulation as well as in tissue-resident cell pools. Tfr play a critical role in controlling the Tfh and B-cells involved in donor-specific humoral immunity.(53, 54) Recent studies have suggested that reduced Tfr-to-Tfh ratios in circulation are a risk factor for allograft dysfunction or failure(97, 98). In our short-term follow up, we found that Tfr-to-Tfh ratios in peripheral blood notably increased after S-CD3e-IT treatment in OVA-immunized mice, but only marginally in nonimmunized mice (Figure 4C). For the tissue-resident (IVS−) compartments, mesenteric LNs showed a notable increase in Tfr-to-Tfh ratios, but the pattern was unclear in the spleen and thymus. Despite these organ-to-organ variations, it is noteworthy that the Tfr-to-Tfh ratios were consistently high in most LN types (except mandibular) in S-CD3e-IT-treated mice (see Figure 5E below). Consistent to these results, previous CD3e-IT-mediated chimerism studies showed an effective and durable modulation of donor-specific humoral responses, while the modulation of donor-specific T-cell immunity was not evident.(34,38)

**Figure 5.**
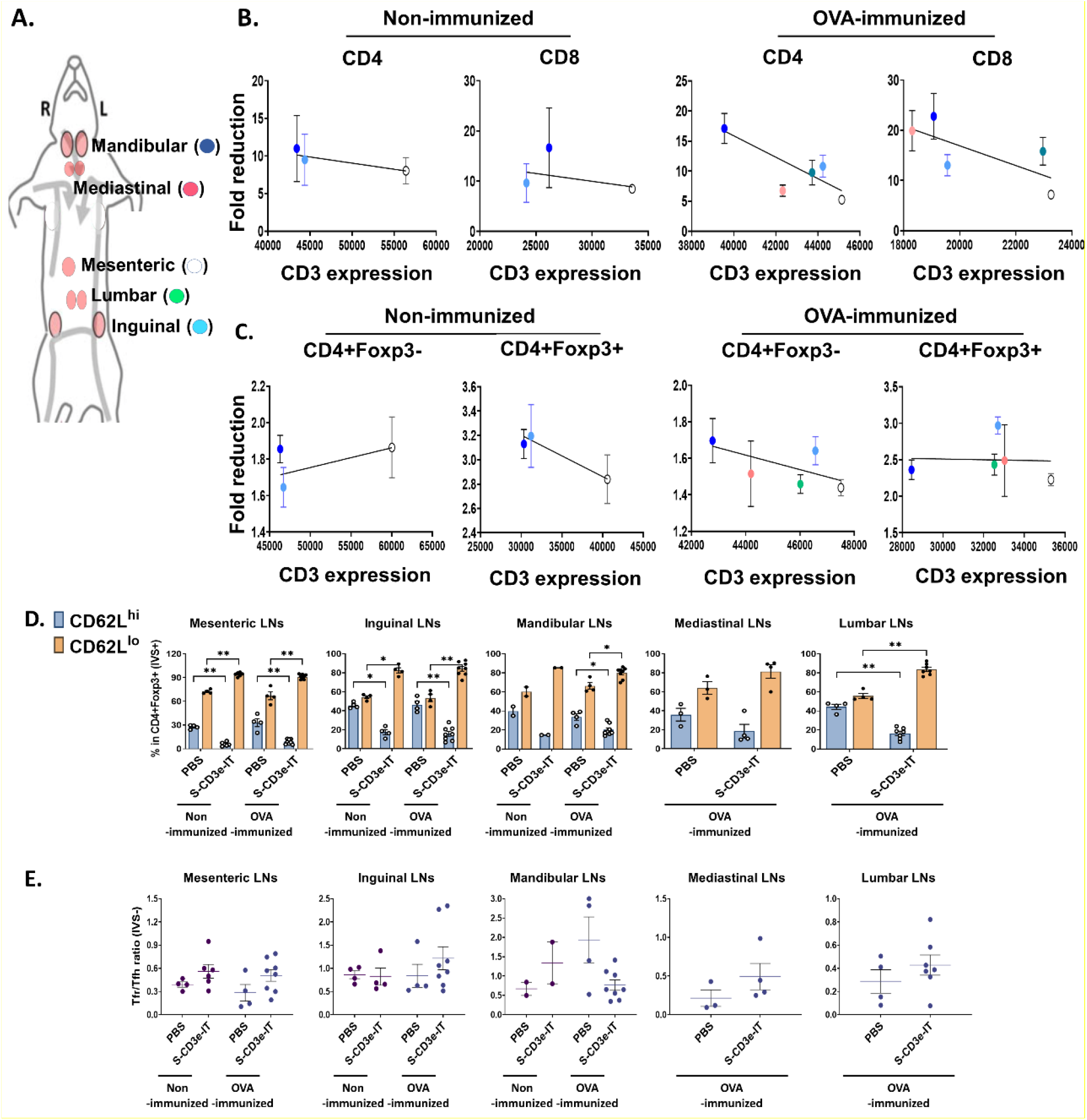
Consistent LN-to-LN variations. (A) Symbols of mesenteric (white), inguinal (light blue), mandibular (blue), mediastinal (red), and lumber (green) LNs are shown. (B) Fold reductions of CD4+ or CD8+ T cells (y-axis; S-CD3e-IT/PBS) and CD3e MFI of these cells in each LNs (x-axis) are shown. There was no positive correlation between CD3e MFI and CD4+ and CD8+ cell depletion rates. (C) Fold reductions of CD4+Foxp3-or CD4+Foxp3+ T cells (y-axis; S-CD3e-IT/PBS) and CD3e MFI of these cells in each LNs (x-axis) are shown. There was no positive correlation between CD3e MFI and CD4+Foxp3-cells. (D) CD62L^lo^ and CD62L^hi^ (% in CD4+Foxp3+ cells) are shown for these LNs. The surviving cells were enriched in CD4+Foxp3+ cell pools with the CD62L^lo^ phenotype. (E) The Tfr-to-Tfh ratios (% in CD4+) are shown for these LNs. S-CD3e-IT-treated mice show higher Tfr-to-Tfh ratios in most of these LNs. Nonimmunized mice were treated with PBS (*n* = 2∼4, depending on LN types) or S-CD3e-IT (*n* = 2∼6). OVA-immunized mice were treated with PBS (*n* = 3∼4) or S-CD3e-IT) (*n* = 4∼8) * *p* < 0.05 and ** *p* < 0.01.

### LN-to-LN variations in T-cell depletion and Treg enrichment

Last, we evaluated LN-to-LN variations in drug efficacy. LNs are major lymphoid organs located in many parts of the body, orchestrating for local Treg-mediated immune regulation in a tissue-specific manner.(52, 72) Studies have shown that drug efficacy in different LNs is highly variable and dependent upon the anatomic sites of the target LNs and drug administration routes, resulting in inconsistent responses to drug regimens.(99–101) Consistent with these reports, we observed considerable LN-to-LN variations in T-cell depletion depending on the type of LNs (Supplemental Figure 3A). We isolated mandibular, inguinal, mesenteric, lumbar, and mediastinal LNs and compared T-cell depletion in these organs (Supplemental Figure 3A). Unlike the other organs tested in this study, there was no correlation between CD3e expression levels and CD4+ or CD8+ T-cell depletion rates among these LN types (Figure 5B). The T-cell depletion rates in these LNs appear to depend on the anatomic positions of these LNs, rather than CD3e expression levels. Mesenteric LN T cells, for example, consistently showed highest CD3e MFI levels for all T-cell subtypes, while mandibular LN T cells consistently showed lowest CD3e MFI among all tested LN types (Figure 5, B and C). Regardless of the differential CD3e expression levels among LNs, however, mandibular LNs consistently showed highest depletion rates for most of T-cell types tested and mesenteric LNs showed lowest depletion rates for the most of T-cell subtypes (Figure 5, A-C). These results are consistent with previous reports addressing the variability of drug efficacy depending on the injection and circulation routes for different LNs.(99–101) Interestingly, however, while the Treg composition (CD62L^hi^ and CD62L^lo^) was notably varied in each type of LN, we found all LN types showed a substantial enrichment of CD3^dim^CD62L^lo^ Tregs, accounting for >80% of the surviving Tregs, after S-CD3e-IT treatment (Figure 5D). Tfr-to-Tfh ratios were notably increased in most LN types, except mandibular LNs (Figure 5E). These results, therefore, suggest additional layers of treatment resistance and the potential need for treatment optimization to control CD3e-IT efficacy in different types of LNs.

## Discussion

Successful application of precision anti-T-cell biologics requires a detailed understanding of both disease etiology and the *in vivo* efficacy of the drugs in use. This study demonstrates an extensive pharmacodynamics analysis of CD3e-IT treatment using a new murine testing system. Our mouse model enabled a detailed evaluation of CD3e-IT-mediated T-cell depletion in multiple organs for both circulating and tissue-resident cells, evaluated separately. Differential CD3e expression among differing T-cell subsets, as well as organ-to-organ variations in CD3e expression, were identified as a main driver for Treg enrichment and the reshaping of organ-specific T-cell composition after CD3e-IT treatment. Differences in CD3e expression on the surface of CD4+ and CD8+ T cells, as well as Tregs are consistent with those of previous murine and human studies.(18, 102) We also found that Tregs, particularly the CD62L^lo^ effector subset, display relatively low levels of CD3e on the cell surface; this difference perhaps provides a survival advantage for CD62L^lo^ effector Tregs. Tregs are also known to have higher amounts of CD3e isoforms with an undegraded N terminal and to be relatively insensitive to antibody stimuli;(19, 20) these factors likely bolster their resistance to CD3e-IT-mediated cell death as well.

Notably, we also found substantial organ-to-organ variations in T-cell depletion. We identified (i) variations in CD3e expression among tissue-resident T cells of different organs as a major factor contributing to the CD3e-IT-mediated reshaping of local T-cell composition. The organ-to-organ variations in CD3e expression may reflect the unique immunologic status of tissue-resident local T cells mostly controlled by local stimuli, maintaining a distinct TCR/CD3 repertoire.(45, 55) Additional factors we identified that contribute to the organ-to-organ variations in T-cell depletion include (ii) the unique functional properties of each organ (e.g. the thymus producing new T cells and the bone marrow actively retaining Tregs), and (iii) the dependency of anatomic locations of LNs in CD3e-IT efficacy which is due likely to the effects of drug injection and circulation routes.(99–101) All these factors (i – iii) may collectively affects CD3e-IT treatment efficacy in local organs and the reshaping of local T-cell immunity following CD3e-IT-treatment.

Mounting evidence points to the critical roles of Tregs in promoting tolerance in various treatment settings.(30–33) Yet Tregs can act like a double-edged sword insofar as it also suppresses anti-cancer T-cell immunity, making the local environment more favorable for cancer growth.(55) Despite the crucial need, Treg enrichment and their functional properties in CD3e-IT treatment remained unclear. Similar to the transient enrichment of Tregs in the early tolerance-induction period of CD3e-mAb (145-2C11) treatment in mice,(18, 73, 82–89) we also found a substantial enrichment of Tregs after S-CD3e-IT treatment. The phenotypic properties of the surviving Tregs in S-CD3e-IT-treated mice were, however, distinct from those in CD3e-mAb-treated mice. Unlike CD3e-mAb – which induces, not kills, Tregs to lead to active tolerance mechanisms involving CD62L^hi^ Tregs (18, 78–82) – CD3e-IT preferentially kills CD3e^hi^ T cells and enriches CD62L^lo^ Tregs with the CD3e^dim^ phenotype. The CD62L^lo^ effector Tregs with CD3e^dim^ phenotype may have a lower tolerogenic potential than CD62L^hi^ Tregs because, first, CD62L^hi^ Tregs, not the CD62L^lo^ counterparts, are reported to be the major population that provides long-lasting immune tolerance,(78–80) and second, CD62L^lo^ effector Tregs require constant CD3 activation to proliferate and maintain their functional properties.(93–95) The surviving Tregs with the CD3e^dim^ CD62L^lo^ phenotype were only marginally activated in CD3e-IT treated mice. Given the critical importance of Treg functions during the early tolerance-induction period, this difference in the Treg phenotype may have an important implication for CD3e-IT-mediated transient chimerism studies.(29, 44) We also observed a notable increase in Tfr-to-Tfh ratios in most LNs following CD3e-IT treatment; this increase may partly explain the robust and durable humoral immune modulation of donor-specific antibody responses in these transient chimerism studies.(29, 44) Treg-mediated donor T-cell tolerance was not evident in swine and monkey models,(29, 44) but longitudinal tracking of the Tregs in long-term induction settings using our new mouse testing model may provide useful means to further investigate these important questions.

CD3e-immunotoxins are a promising short-term, precision treatment option for various therapies that require effective depletion of T cells. Vascular leak syndrome (VLS) has been the major dose-limiting toxicity, but the development of the “second generation” recombinant CD3e-ITs has significantly improved safety profiles in clinical and preclinical studies(1, 14, 16, 103, 104). The recent identification of VLS-inducing (x)D(y) motif of toxin molecules offers new opportunities to further reduce VLS.(104–110) Despite the significant improvement, the treatment efficacy of CD3e-IT has been often limited by unknown causes. The new insights gained in this study into potential treatment resistance mechanisms of CD3e-ITs and mechanisms of Treg enrichment in different organs can serve as a new foundation on which to further improve this promising precision medicine in clinical and preclinical studies.

## Methods

### Mice

Male C57BL/6J mice were obtained from the Jackson Laboratory (Bar Harbor, ME). All mice in this study were maintained by the guidelines of the Ohio State University (OSU). 4∼6 months old male mice were used in this study.

### S-CD3e-IT preparation

To generate murine version of S-CD3e-IT, Fc-silent CD3e monoclonal antibody (S-CD3e-mAb; Absolute Antibody, Upper Heyford, UK) firstly was biotinylated as previously described(46). Anti-murine version of S-CD3e-IT was prepared by conjugating biotinylated-S-CD3e-mAb with streptavidin-ZAP (Advanced Targeting System, San Diego, Calif) in a 1:1 molar ratio as previously described(46).

### Immunization

For immunizing mice, male C57BL/6J mice were received intraperitoneally with ovalbumin (OVA) emulsified in Freund’s complete adjuvant (CFA) followed by OVA emulsified in Freund’s incomplete adjuvant (IFA) one week after the first OVA/CFA injection.

### *In vivo* experiments

Male C57BL/6J mice were injected into retro-orbital sinus with 15 μg S-CD3e-IT in sterile 200 μl PBS twice a day for four consecutive days and were euthanized on day 6 as previously described(46). On the day of euthanasia, mice were injected into the retro-orbital sinus with a total of 3 mg of PE/Cy7-CD45.2 (104, BioLegend) in 200 μl sterile DPBS. After 3 minutes of injection, the mice were euthanized following the previously established protocols(47, 111). The peripheral blood was collected from the heart. The spleen, five LNs (mesenteric, inguinal, mandibular, mediastinal, and lumber LNs), lung, thymus, and bone marrow (right femur) were harvested, and tissues were processed to isolate leukocytes following previous studies with modifications(112, 113).

### Flow cytometry

The collected leukocytes from peripheral blood, spleen, five LNs, lung, thymus, and bone marrow were incubated with Fc-blocking TruStain FcX™ (anti-mouse CD16/32, BioLegend) antibody, followed by Zombie aqua (Live/dead indicator, BioLegend). The cells were stained with fluorescence-labeled antibodies. The following antibodies were purchased from BioLegend: BV650-CD45.2 (104), PE/Cy7-CD45.2 (104), PE/Dazzle594-CD69 (H1.2F3), Alexa Flour 700-CD3e (500A2), Pacific Blue-CD4 (RM4-5), BV570-CD8 (53-6.7), BV711-CD19 (6D5), PerCP/Cy5.5-CD44 (IM7), APC/Cy7-CD62L (MEL-14), Alexa Flour 647-CXCR5 (L138D7), BV421-PD-1 (RMP1-3D), BV785-CD25 (PC61), and BV605-Ki67 (16A8). PE/Cy5-Foxp3 (FJK-16s) was purchased from Invitrogen. All samples were acquired on a Cytek Aurora, and the data were analyzed with FlowJo software (BD Bioscience). The absolute number of leukocytes was calculated using CountBright™ as previously described(46).

### Foxp3 staining

Staining the transcription factor, Foxp3, was carried out using the Foxp3/Transcription Factor Staining buffer set (Invitrogen, Waltham, MA, USA) and according to the recommended manufacturer manual.

### Statistics

Data represent Mean ± SEM. Mann-Whitney test using Prism 9 software (GraphPad Software, La Jolla, CA) was used to analyze if there were significant differences between PBS-treated and S-CD3e-IT-treated mice. P value less than 0.05 was considered significant.

### Study approval

All animal experiments were conducted with the protocol approved by the OSU Animal Care and Use Committee.

## Supporting information

CD3e-IT pharmacodynamics_Supplemental file

## Author Contributions

S.K (Sanggu Kim) and SK (Shihyoung Kim) designed all experiments. S.K(Shihyoung Kim), R.K.S, A.B, H.Y, S.G.C, H.C, S.G, and G.E.L carried out the animal experiment. S.K (Shihyoung Kim), A.B, and S.G.C performed flow cytometry. S.K (Sanggu Kim) and S.K (Shihyoung Kim) analyzed the data. S.K (Sanggu Kim) provided the reagents. S.K (Sanggu Kim) and S.K (Shihyoung Kim) wrote the manuscript. Z.W, C.A.H, J.C.R, and N.P.M.L provided critical appraisal of the manuscript.

## Acknowledgments

This research was funded by the National Institutes of Health (NHLBI R00HL116234, NHGRI R01HG010318, and NHGRI R21HG010108; American Society of Hematology Scholar Award to S.K. (Sanggu Kim); and C. Glenn Barber Fund to S.K. (Shihyoung Kim).

## Notes

### Competing Interest Statement

The authors have declared no competing interest.

## References

1. Frankel AE, et al. Resimmune, an anti-CD3ε recombinant immunotoxin, induces durable remissions in patients with cutaneous T-cell lymphoma. Haematologica 2015;100(6):794–800.

2. Hu H, et al. Depletion of T Lymphocytes with Immunotoxin Retards the Progress of Experimental Allergic Encephalomyelitis in Rhesus Monkeys. Cellular Immunology 1997;177(1):26–34.

3. Jonker M, et al. Long-term kidney graft survival by delayed T cell ablative treatment in rhesus monkeys1,2. Transplantation 2002;73(6):874–880.

4. Leonard DA, et al. Vascularized Composite Allograft Tolerance Across MHC Barriers in a Large Animal Model. American Journal of Transplantation 2014;14(2):343–355.

5. Huang CA, et al. Stable mixed chimerism and tolerance using a nonmyeloablative preparative regimen in a large-animal model. J Clin Invest 2000;105(2):173–181.

6. Wang Z, et al. Development of a Diphtheria Toxin Based Antiporcine CD3 Recombinant Immunotoxin. Bioconjugate Chem. 2011;22(10):2014–2020.

7. Schwarze ML, et al. Mixed hematopoietic chimerism induces long-term tolerance to cardiac allografts in miniature swine. The Annals of Thoracic Surgery 2000;70(1):131–138.

8. Thompson J, et al. Improved binding of a bivalent single-chain immunotoxin results in increased efficacy for in vivo T-cell depletion. Protein Eng Des Sel 2001;14(12):1035–1041.

9. Frankel AE, et al. Resimmune, an anti-CD3 recombinant immunotoxin, induces durable remissions in patients with cutaneous T-cell lymphoma. Haematologica 2015;100(6):794–800.

10. Thompson J, et al. An Anti-CD3 Single-chain Immunotoxin with a Truncated Diphtheria Toxin Avoids Inhibition by Pre-existing Antibodies in Human Blood. Journal of Biological Chemistry 1995;270(47):28037–28041.

11. Johnson V e., et al. Genetic markers associated with progression in early mycosis fungoides. Journal of the European Academy of Dermatology and Venereology 2014;28(11):1431–1435.

12. Vaqué JP, et al. PLCG1 mutations in cutaneous T-cell lymphomas. Blood 2014;123(13):2034– 2043.

13. Kim J-S, Jun S-Y, Kim Y-S. Critical Issues in the Development of Immunotoxins for Anticancer Therapy. Journal of Pharmaceutical Sciences 2020;109(1):104–115.

14. Woo JH, et al. Preclinical studies in rats and squirrel monkeys for safety evaluation of the bivalent anti-human T cell immunotoxin, A-dmDT390–bisFv(UCHT1). Cancer Immunol Immunother 2008;57(8):1225–1239.

15. Hamawy MM, et al. Activation of T lymphocytes for adhesion and cytokine expression by toxin-conjugated anti-CD3 monoclonal antibodies. Transplantation 1999;68(5):693–698.

16. Matar AJ, et al. Effect of pre-existing anti-diphtheria toxin antibodies on T cell depletion levels following diphtheria toxin-based recombinant anti-monkey CD3 immunotoxin treatment. Transpl Immunol 2012;27(1):52–54.

17. You S. Differential Sensitivity of Regulatory and Effector T Cells to Cell Death: A Prerequisite for Transplant Tolerance. Frontiers in Immunology 2015;6:242.

18. Valle A, et al. Heterogeneous CD3 Expression Levels in Differing T Cell Subsets Correlate with the In Vivo Anti-CD3–Mediated T Cell Modulation. J.I. 2015;194(5):2117–2127.

19. Criado G, et al. Variability of invariant mouse CD3ϵ chains detected by anti-CD3 antibodies. European Journal of Immunology 2000;30(5):1469–1479.

20. Rojo JM, et al. Characteristics of TCR/CD3 complex CD3? chains of regulatory CD4+ T (Treg) lymphocytes: role in Treg differentiation in vitro and impact on Treg in vivo. Journal of Leukocyte Biology 2014;95(3):441–450.

21. Liu YY, et al. Polymorphisms of CD3ε in cynomolgus and rhesus monkeys and their relevance to anti-CD3 antibodies and immunotoxins. Immunology & Cell Biology 2007;85(5):357–362.

22. Dieffenbach M, Pastan I. Mechanisms of Resistance to Immunotoxins Containing Pseudomonas Exotoxin A in Cancer Therapy. Biomolecules 2020;10(7):979.

23. Neville DMJ, et al. A New Reagent for the Induction of T-Cell Depletion, Anti-CD3-CRM9. Journal of Immunotherapy 1996;19(2):85–92.

24. Fechner JH, et al. Mechanisms of Tolerance Induced by an lmmunotoxin Against CD∼E in a Rhesus Kidney Allograft Model:1.

25. Huang CA, et al. IN VIVO T CELL DEPLETION IN MINIATURE SWINE USING THE SWINE CD3 IMMUNOTOXIN, pCD3-CRM91. Transplantation 1999;68(6):855–860.

26. Fuchimoto Y, et al. Mixed chimerism and tolerance without whole body irradiation in a large animal model. J Clin Invest 2000;105(12):1779–1789.

27. Cina RA, et al. Stable Multilineage Chimerism without Graft versus Host Disease Following Nonmyeloablative Haploidentical Hematopoietic Cell Transplantation. Transplantation 2006;81(12):1677–1685.

28. Wang Z, et al. Improvement of a Recombinant Anti-Monkey Anti-CD3 Diphtheria Toxin Based Immunotoxin by Yeast Display Affinity Maturation of the scFv. Bioconjugate Chem. 2007;18(3):947–955.

29. Pathiraja V, et al. Tolerance of Vascularized Islet-Kidney Transplants in Rhesus Monkeys. Am J Transplant 2017;17(1):91–102.

30. Pathak S, Meyer EH. Tregs and Mixed Chimerism as Approaches for Tolerance Induction in Islet Transplantation. Frontiers in Immunology 2021;11. doi:10.3389/fimmu.2020.612737

31. Sayitoglu EC, Freeborn RA, Roncarolo MG. The Yin and Yang of Type 1 Regulatory T Cells: From Discovery to Clinical Application. Frontiers in Immunology 2021;12. doi:10.3389/fimmu.2021.693105

32. Ruiz P, et al. Alloreactive Regulatory T Cells Allow the Generation of Mixed Chimerism and Transplant Tolerance. Frontiers in Immunology 2015;6. doi:10.3389/fimmu.2015.00596

33. Joffre O, et al. Prevention of acute and chronic allograft rejection with CD4+CD25+Foxp3+ regulatory T lymphocytes. Nat Med 2008;14(1):88–92.

34. Wamala I, et al. Recombinant anti-monkey CD3 immunotoxin depletes peripheral lymph node T lymphocytes more effectively than rabbit anti-thymocyte globulin in naïve baboons. Transplant Immunology 2013;29(1–4):60–63.

35. Watanabe R, et al. Human skin is protected by four functionally and phenotypically discrete populations of resident and recirculating memory T cells. Sci Transl Med 2015;7(279):279ra39.

36. Knechtle SJ, et al. Primate renal transplants using immunotoxin 1998;124(2):10.

37. Knechtle SJ, et al. FN18-CRM9 IMMUNOTOXIN PROMOTES TOLERANCE IN PRIMATE RENAL ALLOGRAFTS1. Transplantation 1997;63(1):1–6.

38. Torrealba JR, et al. Metastable Tolerance to Rhesus Monkey Renal Transplants Is Correlated with Allograft TGF-β1 + CD4 + T Regulatory Cell Infiltrates. J Immunol 2004;172(9):5753– 5764.

39. Torrealba JR, et al. Immunotoxin-treated rhesus monkeys: a model for renal allograft chronic rejection1. Transplantation 2003;76(3):524–530.

40. Page EK, et al. Enhanced De Novo Alloantibody and Antibody-Mediated Injury in Rhesus Macaques. American Journal of Transplantation 2012;12(9):2395–2405.

41. Gupta PK, et al. The pursuit of transplantation tolerance: new mechanistic insights. Cell Mol Immunol 2019;16(4):324–333.

42. Graca L, et al. Donor-specific transplantation tolerance: The paradoxical behavior of CD4+CD25+ T cells. Proceedings of the National Academy of Sciences 2004;101(27):10122– 10126.

43. Zuber J, Sykes M. Mechanisms of Mixed Chimerism-Based Transplant Tolerance. Trends in Immunology 2017;38(11):829–843.

44. Duran-Struuck R, et al. Lack of Antidonor Alloantibody Does Not Indicate Lack of Immune Sensitization: Studies of Graft Loss in a Haploidentical Hematopoietic Cell Transplantation Swine Model. Biol Blood Marrow Transplant 2012;18(11):1629–1637.

45. Masopust D, Soerens AG. Tissue-Resident T Cells and Other Resident Leukocytes. Annu Rev Immunol 2019;37:521–546.

46. Kim S, et al. Comparison of CD3e Antibody and CD3e-sZAP Immunotoxin Treatment in Mice Identifies sZAP as the Main Driver of Vascular Leakage. Biomedicines 2022;10(6):1221.

47. Anderson KG, et al. Intravascular staining for discrimination of vascular and tissue leukocytes. Nat Protoc 2014;9(1):209–222.

48. Potter EL, et al. Measurement of leukocyte trafficking kinetics in macaques by serial intravascular staining. Science Translational Medicine 2021;13(576):eabb4582.

49. Ochando JC, et al. Lymph Node Occupancy Is Required for the Peripheral Development of Alloantigen-Specific Foxp3 + Regulatory T Cells. J Immunol 2005;174(11):6993–7005.

50. Schneider MA, et al. CCR7 is required for the in vivo function of CD4+ CD25+ regulatory T cells. Journal of Experimental Medicine 2007;204(4):735–745.

51. Zhang N, et al. Regulatory T Cells Sequentially Migrate from Inflamed Tissues to Draining Lymph Nodes to Suppress the Alloimmune Response. Immunity 2009;30(3):458–469.

52. Lathrop SK, et al. Antigen-specific peripheral shaping of the natural regulatory T cell population. Journal of Experimental Medicine 2008;205(13):3105–3117.

53. Vaeth M, et al. Follicular regulatory T cells control humoral autoimmunity via NFAT2-regulated CXCR5 expression. Journal of Experimental Medicine 2014;211(3):545–561.

54. Linterman MA, et al. Foxp3+ follicular regulatory T cells control the germinal center response. Nat Med 2011;17(8):975–982.

55. Wei S, Kryczek I, Zou W. Regulatory T-cell compartmentalization and trafficking. Blood 2006;108(2):426–431.

56. Mathes DW, et al. Tolerance to Vascularized Composite Allografts in Canine Mixed Hematopoietic Chimeras. Transplantation 2011;92(12):1301–1308.

57. Eljaafari A, et al. Isolation of Regulatory T Cells in the Skin of a Human Hand-Allograft, Up to Six Years Posttransplantation. Transplantation 2006;82(12):1764–1768.

58. Vallera DA, et al. Molecular modification of a recombinant anti-CD3ε-directed immunotoxin by inducing terminal cysteine bridging enhances anti-GVHD efficacy and reduces organ toxicity in a lethal murine model. Blood 2000;96(3):1157–1165.

59. Vallera D, et al. Therapy for ongoing graft-versus-host disease induced across the major or minor histocompatibility barrier in mice with anti-CD3F(ab’)2-ricin toxin A chain immunotoxin. Blood 1995;86(11):4367–4375.

60. Kim G-B, et al. A fold-back single-chain diabody format enhances the bioactivity of an antimonkey CD3 recombinant diphtheria toxin-based immunotoxin. Protein Engineering Design and Selection 2007;20(9):425–432.

61. Vallera DA, Panoskaltsis-Mortari A, Blazar BR. Renal dysfunction accounts for the dose limiting toxicity of DT390anti-CD3sFv, a potential new recombinant anti-GVHD immunotoxin. Protein Engineering Design and Selection 1997;10(9):1071–1076.

62. Keijzer C, et al. Treg Inducing Adjuvants for Therapeutic Vaccination Against Chronic Inflammatory Diseases. Frontiers in Immunology 2013;4(4). doi:10.3389/fimmu.2013.00245

63. Wu T, et al. Immunosuppressive drugs on inducing Ag-specific CD4+CD25+Foxp3+ Treg cells during immune response in vivo. Transplant Immunology 2012;27(1):30–38.

64. Riteau N, et al. Water-in-Oil–Only Adjuvants Selectively Promote T Follicular Helper Cell Polarization through a Type I IFN and IL-6–Dependent Pathway. J.I. 2016;197(10):3884– 3893.

65. Beura LK, et al. T Cells in Nonlymphoid Tissues Give Rise to Lymph-Node-Resident Memory T Cells. Immunity 2018;48(2):327-338.e5.

66. Beura LK, et al. Normalizing the environment recapitulates adult human immune traits in laboratory mice. Nature 2016;532(7600):512–516.

67. Shevach EM. CD4+CD25+ suppressor T cells: more questions than answers. Nat Rev Immunol 2002;2(6):389–400.

68. Sakaguchi S. Naturally arising Foxp3-expressing CD25+CD4+ regulatory T cells in immunological tolerance to self and non-self. Nat Immunol 2005;6(4):345–352.

69. Zou L, et al. Bone Marrow Is a Reservoir for CD4 + CD25 + Regulatory T Cells that Traffic through CXCL12/CXCR4 Signals. Cancer Res 2004;64(22):8451–8455.

70. Stephens LA, et al. Human CD4+CD25+ thymocytes and peripheral T cells have immune suppressive activity in vitro. European Journal of Immunology 2001;31(4):1247–1254.

71. Curiel TJ, et al. Specific recruitment of regulatory T cells in ovarian carcinoma fosters immune privilege and predicts reduced survival. Nat Med 2004;10(9):942–949.

72. Huang M-T, et al. Lymph node trafficking of regulatory T cells is prerequisite for immune suppression. Journal of Leukocyte Biology 2016;99(4):561–568.

73. Mehta DS, et al. Partial and transient modulation of the CD3-T-cell receptor complex, elicited by low-dose regimens of monoclonal anti-CD3, is sufficient to induce disease remission in non-obese diabetic mice. Immunology 2010;130(1):103–113.

74. Yamaizumi M, et al. One molecule of diphtheria toxin fragment a introduced into a cell can kill the cell. Cell 1978;15(1):245–250.

75. Campbell DJ, Koch MA. Phenotypical and functional specialization of FOXP3+ regulatory T cells. Nat Rev Immunol 2011;11(2):119–130.

76. Fisson S, et al. Continuous Activation of Autoreactive CD4+ CD25+ Regulatory T Cells in the Steady State. J Exp Med 2003;198(5):737–746.

77. Thornton AM, Shevach EM. Suppressor Effector Function of CD4 + CD25 + Immunoregulatory T Cells Is Antigen Nonspecific. J Immunol 2000;164(1):183–190.

78. Szanya V, et al. The Subpopulation of CD4 + CD25 + Splenocytes That Delays Adoptive Transfer of Diabetes Expresses L-Selectin and High Levels of CCR7. J Immunol 2002;169(5):2461–2465.

79. Ermann J, et al. Only the CD62L+ subpopulation of CD4+CD25+ regulatory T cells protects from lethal acute GVHD. Blood 2005;105(5):2220–2226.

80. Taylor PA, et al. L-Selectinhi but not the L-selectinlo CD4+25+ T-regulatory cells are potent inhibitors of GVHD and BM graft rejection. Blood 2004;104(12):3804–3812.

81. Yang W, et al. Perturbed Homeostasis of Peripheral T Cells Elicits Decreased Susceptibility to Anti-CD3-Induced Apoptosis in Prediabetic Nonobese Diabetic Mice. J Immunol 2004;173(7):4407–4416.

82. Kohm AP, et al. Treatment with Nonmitogenic Anti-CD3 Monoclonal Antibody Induces CD4 + T Cell Unresponsiveness and Functional Reversal of Established Experimental Autoimmune Encephalomyelitis. J Immunol 2005;174(8):4525–4534.

83. von Herrath MG, et al. Nonmitogenic CD3 Antibody Reverses Virally Induced (Rat Insulin Promoter-Lymphocytic Choriomeningitis Virus) Autoimmune Diabetes Without Impeding Viral Clearance. J Immunol 2002;168(2):933–941.

84. Valle A, et al. Rapamycin Prevents and Breaks the Anti-CD3–Induced Tolerance in NOD Mice. Diabetes 2009;58(4):875–881.

85. Belghith M, et al. TGF-β-dependent mechanisms mediate restoration of self-tolerance induced by antibodies to CD3 in overt autoimmune diabetes. Nat Med 2003;9(9):1202–1208.

86. Besançon A, et al. The Induction and Maintenance of Transplant Tolerance Engages Both Regulatory and Anergic CD4+ T cells. Front Immunol 2017;8:218.

87. Penaranda C, Tang Q, Bluestone JA. Anti-CD3 Therapy Promotes Tolerance by Selectively Depleting Pathogenic Cells while Preserving Regulatory T Cells. J.I. 2011;187(4):2015–2022.

88. Peng B, et al. Anti-CD3 antibodies modulate anti–factor VIII immune responses in hemophilia A mice after factor VIII plasmid-mediated gene therapy. Blood 2009;114(20):4373–4382.

89. Cook DP, et al. Intestinal Delivery of Proinsulin and IL-10 via Lactococcus lactis Combined With Low-Dose Anti-CD3 Restores Tolerance Outside the Window of Acute Type 1 Diabetes Diagnosis. Front Immunol 2020;11:1103.

90. Malek TR, et al. CD4 Regulatory T Cells Prevent Lethal Autoimmunity in IL-2R?-Deficient Mice: Implications for the Nonredundant Function of IL-2. Immunity 2002;17(2):167–178.

91. Almeida ARM, et al. Homeostasis of Peripheral CD4 + T Cells: IL-2Rα and IL-2 Shape a Population of Regulatory Cells That Controls CD4 + T Cell Numbers. J Immunol 2002;169(9):4850–4860.

92. Zhang H, et al. Lymphopenia and interleukin-2 therapy alter homeostasis of CD4+CD25+ regulatory T cells. Nat Med 2005;11(11):1238–1243.

93. Vahl JC, et al. Continuous T Cell Receptor Signals Maintain a Functional Regulatory T Cell Pool. Immunity 2014;41(5):722–736.

94. Levine AG, et al. Continuous requirement for the TCR in regulatory T cell function. Nat Immunol 2014;15(11):1070–1078.

95. Schmidt AM, et al. Regulatory T Cells Require TCR Signaling for Their Suppressive Function. J.I. 2015;194(9):4362–4370.

96. Page E, et al. Lymphodepletional Strategies in Transplantation. Cold Spring Harb Perspect Med 2013;3(7):a015511.

97. Yan L, et al. Increased circulating Tfh to Tfr ratio in chronic renal allograft dysfunction: a pilot study. BMC Immunol 2019;20(1):26.

98. Niu Q, et al. Immunosuppression Has Long-Lasting Effects on Circulating Follicular Regulatory T Cells in Kidney Transplant Recipients. Front Immunol 2020;11:1972.

99. Allen TM, Hansen CB, Guo LSS. Subcutaneous administration of liposomes: a comparison with the intravenous and intraperitoneal routes of injection. Biochimica et Biophysica Acta (BBA) - Biomembranes 1993;1150(1):9–16.

100. Pitorre M, et al. Passive and specific targeting of lymph nodes: the influence of the administration route. European Journal of Nanomedicine 2015;7(2):121–128.

101. Thompson CG, et al. Mass Spectrometry Imaging Reveals Heterogeneous Efavirenz Distribution within Putative HIV Reservoirs. Antimicrob. Agents Chemother. 2015;59(5):2944–2948.

102. El Hentati F-Z, et al. Variability of CD3 membrane expression and T cell activation capacity. Cytometry Part B: Clinical Cytometry 2010;78B(2):105–114.

103. Darvishi B, et al. Probable Mechanisms Involved in Immunotoxin Mediated Capillary Leak Syndrome (CLS) and Recently Developed Countering Strategies.. Current molecular medicine [published online ahead of print: 2018]; doi:10.2174/1566524018666181004120112

104. Smallshaw JE, et al. Genetic engineering of an immunotoxin to eliminate pulmonary vascular leak in mice. Nat Biotechnol 2003;21(4):387–391.

105. Weldon JE, et al. A protease-resistant immunotoxin against CD22 with greatly increased activity against CLL and diminished animal toxicity. Blood 2009;113(16):3792–3800.

106. Weldon JE, et al. A Recombinant Immunotoxin against the Tumor-Associated Antigen Mesothelin Reengineered for High Activity, Low Off-Target Toxicity, and Reduced Antigenicity. Mol Cancer Ther 2013;12(1):48–57.

107. Wang H, et al. Treatment of hepatocellular carcinoma in a mouse xenograft model with an immunotoxin which is engineered to eliminate vascular leak syndrome. Cancer Immunol Immunother 2007;56(11):1775–1783.

108. Baluna R, et al. Evidence for a structural motif in toxins and interleukin-2 that may be responsible for binding to endothelial cells and initiating vascular leak syndrome. Proceedings of the National Academy of Sciences 1999;96(7):3957–3962.

109. Díaz R, et al. Selective CXCR4+ Cancer Cell Targeting and Potent Antineoplastic Effect by a Nanostructured Version of Recombinant Ricin. Small 2018;14(26):e1800665.

110. Cheung LS, et al. Second-generation IL-2 receptor-targeted diphtheria fusion toxin exhibits antitumor activity and synergy with anti–PD-1 in melanoma. Proc Natl Acad Sci U S A 2019;116(8):3100–3105.

111. Anderson K, et al. Cutting Edge: Intravascular Staining Redefines Lung CD8 T Cell Responses. The Journal of Immunology [published online ahead of print: 2012]; doi:10.4049/jimmunol.1201682

112. Yu Y-RA, et al. A Protocol for the Comprehensive Flow Cytometric Analysis of Immune Cells in Normal and Inflamed Murine Non-Lymphoid Tissues. PLoS One 2016;11(3):e0150606.

113. Misharin AV, et al. Flow Cytometric Analysis of Macrophages and Dendritic Cell Subsets in the Mouse Lung. Am J Respir Cell Mol Biol 2013;49(4):503–510.

