## Supplementary material for "CD3e-immunotoxin spares CD62L^lo^ Tregs and reshapes organ-specific T-cell composition by preferentially depleting CD3e^hi^ T cells": CD3e-IT pharmacodynamics_Supplemental file

**Supplemental Data**

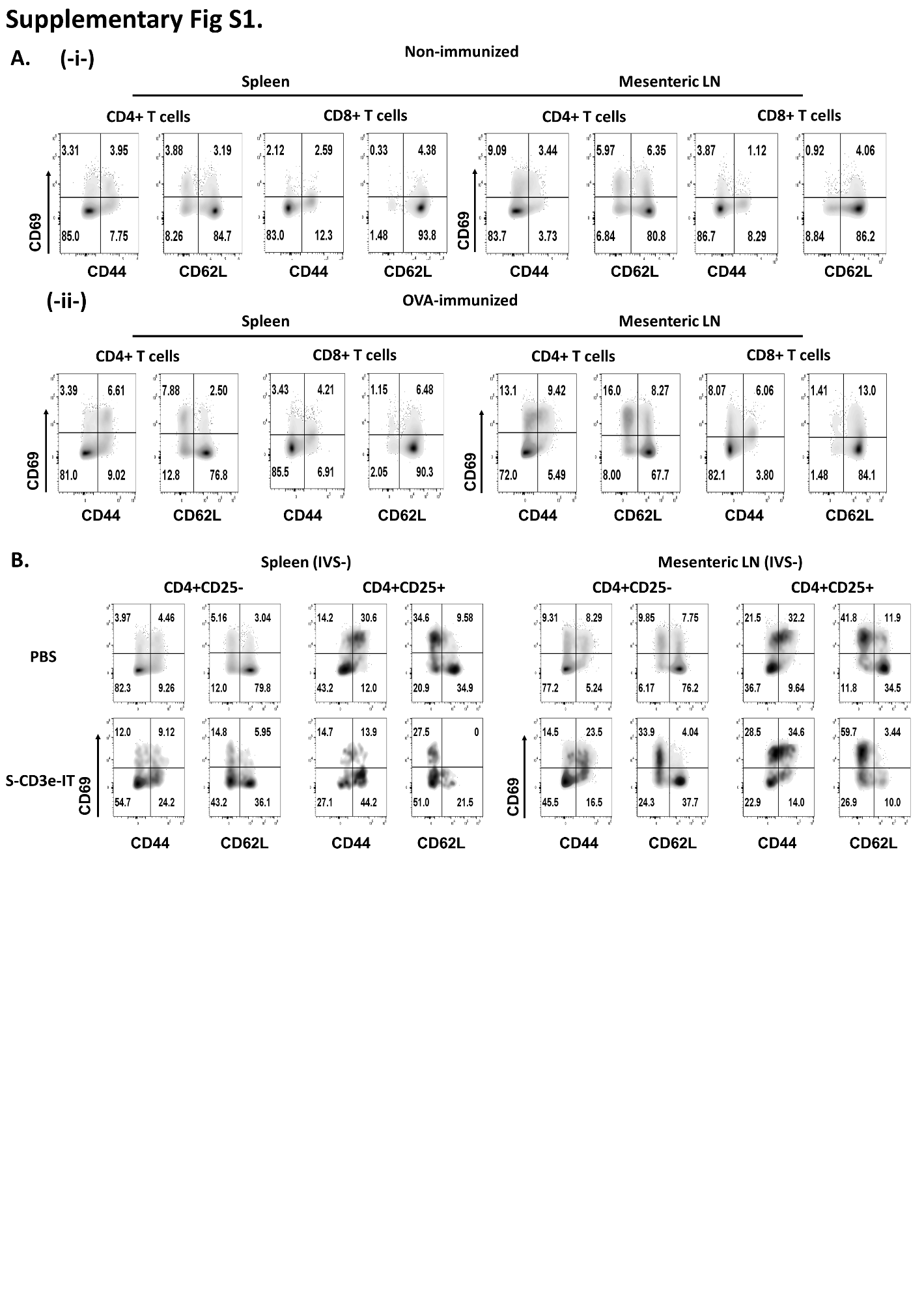

Supplementary Figure 1. (A) Flow image shows the change of CD69+ cells before (i) and after OVA immunization (ii) in spleen and mesenteric LN. The surface CD69 expression (y-axis) is compared on CD44 or CD62L expression (x-axis) for CD4+ T cells (left) and CD8+ T cells (right). CD69 expression was notably increased in LNs cells of OVA-immunized mice compared to non-immunized mice. (B) The percentage change of CD69 expression in CD4+CD25- and CD4+CD25- cells in tissue-resident (IVS-) spleen (left four panels) and mesenteric LN (right four panels) after S-CD3e-IT treatment. PBS treatment (upper panel) and S-CD3-IT (lower panel) are compared.

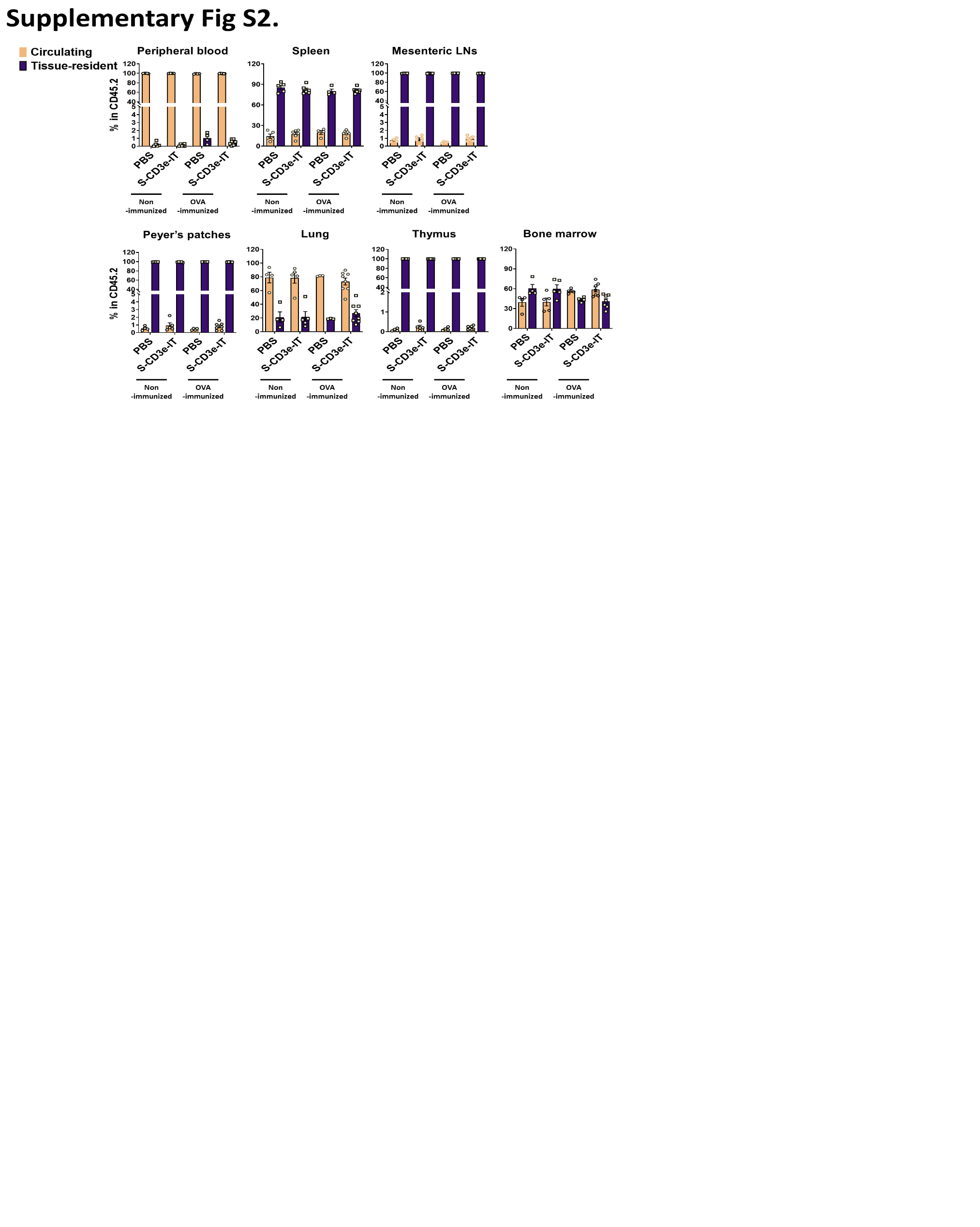

Supplementary Figure 2. The percentage of circulating (IVS+) and tissue-resident (IVS-) cells was shown for peripheral blood, Spleen, mesenteric LNs, Peyer's patches, lung, thymus, and bone marrow. Non-immunized mice (PBS; *n* = 4~5 depending on tissues), non-immunized mice (S-CD3e-IT; *n* = 4~6), OVA-immunized mice(PBS; *n* = 3~4), and OVA-immunized mice (S-CD3e-IT; *n* = 7~8), were compared.

d
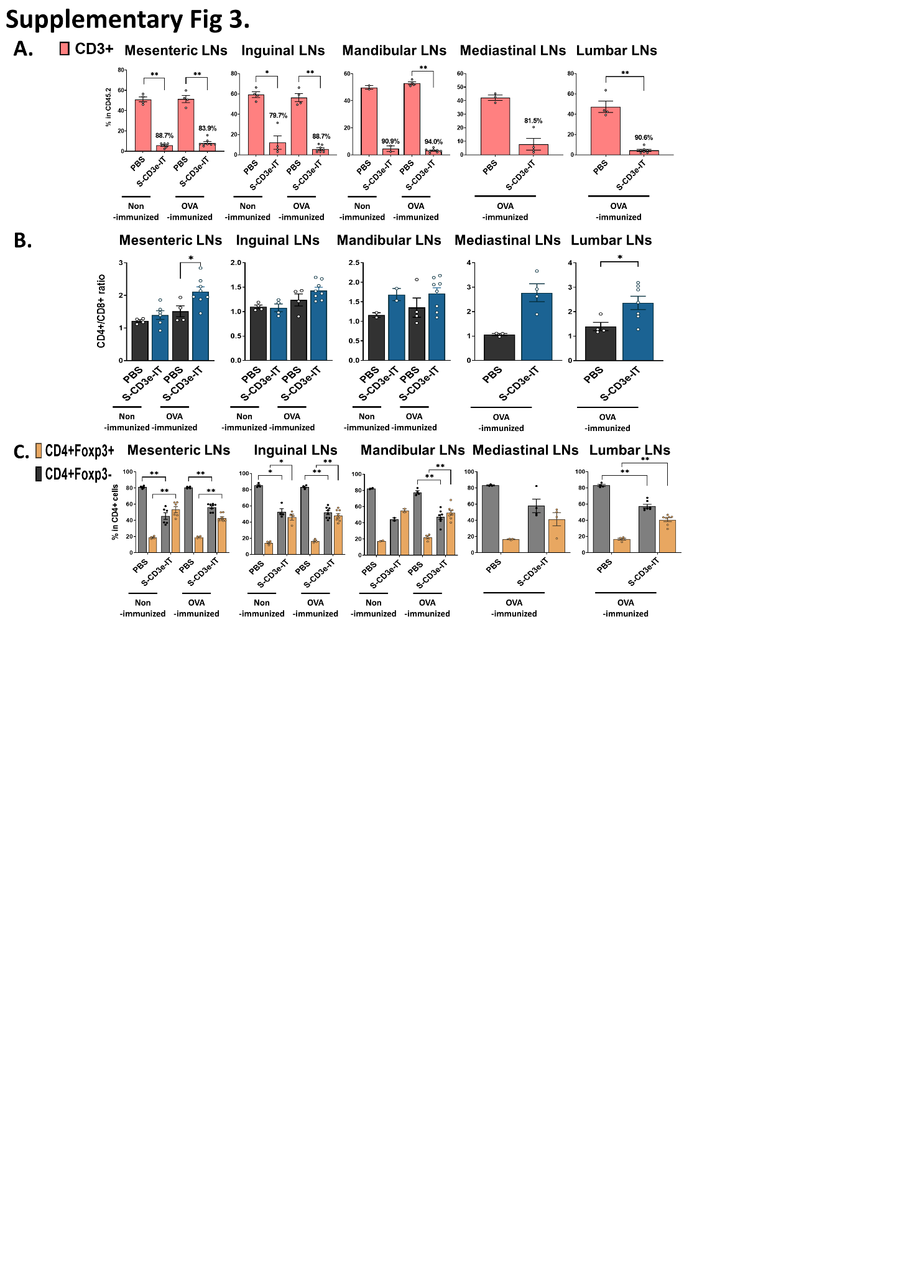

Supplementary Figure 3. (A) CD3+ T cells are shown (y-axis; % in CD45.2+ leukocytes) for the five LNs (mesenteric, inguinal, mandibular, mediastinal, and lumbar LNs) in different anatomic sites. (B) The ratio of CD4+ to CD8+ is shown for these LNs. The ratio increased following S-CD3e-IT treatment. (C) CD4+Foxp3- and CD4+Foxp3+ (% in CD4+ cells) are shown for these LNs. Non-immunized mice (PBS; *n* = 2~4 depending on tissues) non-immunized mice (S-CD3e-IT; *n* = 2~6), OVA-immunized mice (PBS; *n* = 3~4), and OVA-immunized mice (S-CD3e-IT; *n* = 4~8), were compared. (* *p* < 0.05 and ** *p* < 0.01).

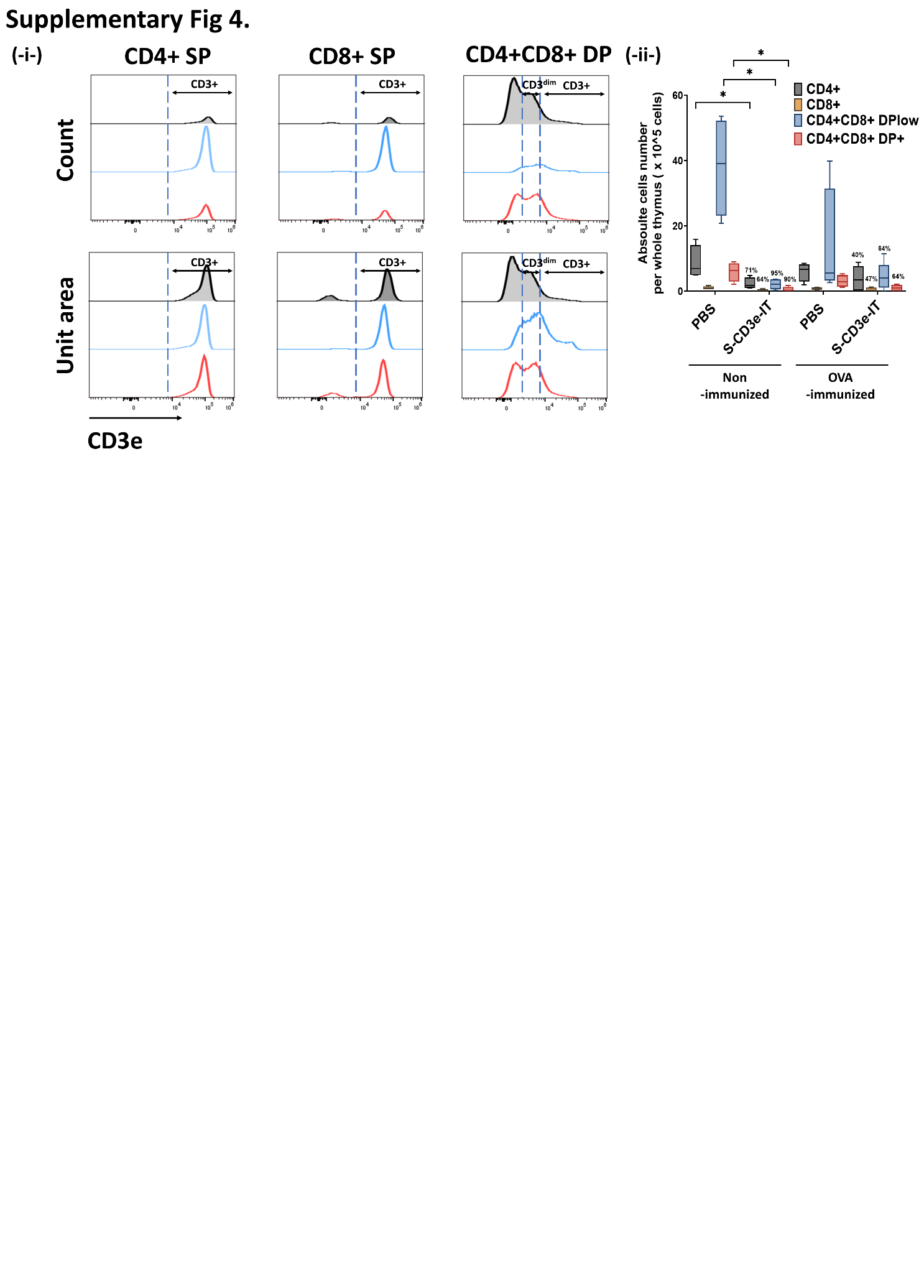

Supplementary Figure 4. **(-i-)** Mean fluorescent intensity (MFI) of CD3e on CD4+, CD8+ single positive (SP), and CD4+CD8+ double positive (DP) cells (**x-axis**) and cell count (**y-axis, upper panel**) or unit area (**y-axis, lower panel**) are shown for the thymus. **(-ii-)** Absolute cell count (per the thymus) for CD4+CD3+, CD8+CD3+ SP, CD4+CD8+CD3^lo^, and CD4+CD8+CD3+ cells are shown for the thymus. Cell numbers were calculated based on CountBright Absolute counting beads and the total cell counts per tissue. Non-immunized mice (PBS; *n* = 4 depending on tissues) non-immunized mice (S-CD3e-IT; *n* = 5), OVA-immunized mice (PBS; *n* = 4), and OVA-immunized mice (S-CD3e-IT; *n* = 7), were compared. (* *p* < 0.05).

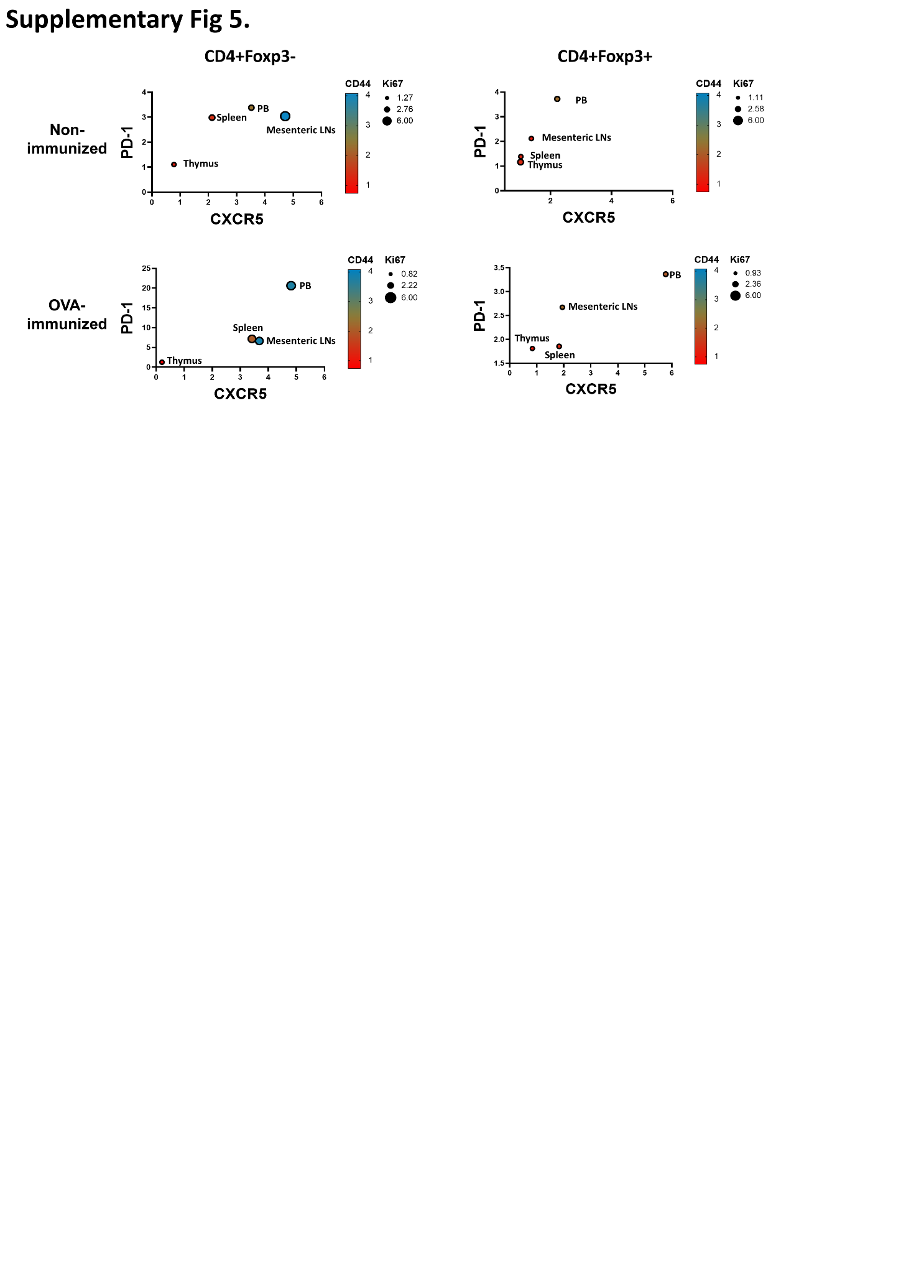

Supplementary Figure 5. The fold change (S-CD3e-IT / PBS) of CXCR5+ (x-axis), PD-1+ (y-axis), CD44+ (color scheme from red to blue), and Ki67+ markers (circle size) for CD4+Foxp3- (left panels) and CD4+Foxp3+ cells (right panels) are shown for peripheral blood, spleen, mesenteric LN, and thymus. Nonimmunized mice were treated with PBS (*n* = 4~5, depending on organs) or S-CD3e-IT (*n* = 4~6). OVA-immunized mice were treated with PBS (*n* = 3~4) or S-CD3e-IT (*n* = 7~8).
